# Ultrasound-guided Photoacoustic image Annotation Toolkit in MATLAB (PHANTOM) for preclinical applications

**DOI:** 10.1101/2023.11.07.565885

**Authors:** Allison Sweeney, Aayush Arora, Skye Edwards, Srivalleesha Mallidi

**Affiliations:** Department of Biomedical Engineering, Tufts University, Medford, MA, United States; Wellman Center for Photomedicine, Massachusetts General Hospital, Boston, MA, United States

## Abstract

Depth-dependent fluence-compensation in photoacoustic (PA) imaging is paramount for accurate quantification of chromophores from deep tissues. Here we present a user-friendly toolkit named PHANTOM (PHotoacoustic ANnotation TOolkit for MATLAB) that includes a graphical interface and assists in the segmentation of ultrasound-guided PA images. We modelled the light source configuration with Monte Carlo eXtreme and utilized 3D segmented tissues from ultrasound to generate fluence maps to depth compensate PA images. The methodology was used to analyze PA images of phantoms with varying blood oxygenation and results were validated with oxygen electrode measurements. Two preclinical models, a subcutaneous tumor and a calcified placenta, were imaged and fluence-compensated using the PHANTOM toolkit and the results were verified with immunohistochemistry. The PHANTOM toolkit provides scripts and auxiliary functions to enable biomedical researchers not specialized in optical imaging to apply fluence correction to PA images, enhancing accessibility of quantitative PAI for researchers in various fields.

## 1. Introduction

Photoacoustic imaging (PAI) is a rapidly evolving imaging method in both the clinical and preclinical realm due to its ability to provide high contrast and resolution at deeper penetration depths than other optical imaging modalities[1, 2]. PAI is intrinsically sensitive to the functional and molecular information of biological tissues due to its optical absorption-based contrast mechanism[3, 4]. PAI exhibits versatile applications across diverse disease spaces, manifesting its clinical efficacy in distinct contexts[5, 6]. Endogenous contrast PAI offers the capability to visualize the structure, oxygenation, flow, destruction, and regeneration of blood vessels making it well adapted to the assessment of tumors, stroke, rheumatoid arthritis, placental dysfunction, and other conditions that exhibit irregularities in vascular morphology and function[7–10]. This also allows PAI to serve as a reliable indicator of successful vascular ablation through modalities like photodynamic therapy, photothermal therapy, and sonodynamic therapy [11–14].

PAI signal is generated when nanosecond pulsed light is absorbed by chromophores to cause thermoelastic expansion and contraction, resulting in the generation of transient broadband acoustic waves[15]. PAI can be transparently integrated with traditional ultrasound imaging systems as they utilize the same receiver electronics[16]. There are many benefits of ultrasound-guided photoacoustic imaging (US-PAI) over PAI alone, the most important of which is the additional anatomical context provided by US[17, 18].The amplitude of the photoacoustic signal generated is dependent on the optical absorption properties of the tissue, inherent thermal properties and the light fluence[2, 3, 18]. Specifically, fluence, or the amount of light energy per area delivered to a region of tissue, plays a critical role in the quantification of the PA signal from deeper tissues[19]. Tissue optical properties including absorption and scattering, give rise to heterogeneous fluence distribution in biological samples[20, 21]. As a consequence, PA image quality is hampered by the effects of depth and wavelength dependent attenuation of optical fluence, in which less light energy is delivered to deeper structures which therefore display weaker photoacoustic signals[22, 23]. The effect is known as spectral coloring and is a well-known confounding factor that compromises the accuracy of blood oxygen saturation (StO_2_) measurements[22, 24–26]. StO_2_ has been shown to be a valuable prognostic marker for several clinical and pre-clinical applications and ensuring the accuracy of this parameter will aid in promoting the clinical translation of PAI as an imaging modality[27–36].

Several methodologies were developed to overcome the shortcoming of spectral coloring, with the primary method being fluence compensation approaches that attempt to normalize the distribution of fluence across the image area[19]. The optical transport model method[37, 38], acoustic spectrum-based method[39, 40], ultrasonic tagging of light[41, 42], fluence matching techniques[43], diffuse optical tomography (DOT) enhanced method[23, 44], signal to noise ratio local fluence correction[45], eigenspectral multispectral model[46], learned spectral decoloring[47], and deep learning methods[48–50] have been proposed and implemented. However, the versatile and customizable nature of Monte Carlo (MC) simulations of light propagation makes it a gold-standard method for fluence compensation of photoacoustic images despite many other alternative methods being developed. Although MC algorithms are regarded as having superior accuracy compared to any analytical diffuse light transport model[51], it is still essential to verify them to ensure they contain no numerical errors or unintended biases. The diffusion approximation is the most common computational approach used to solve the radiative transfer equation (RTE)[52–54]. The MC toolkit used in this work, MCX, has been rigorously validated against the diffusion approximation[55] and has been used in fluence compensation and optical property calculation in various techniques[56–60].

A growing amount of pre-clinical work being published in the field of PAI disregards the effect of light attenuation and presumes that derived parameters are accurate[61–77]. As the PAI modality is becoming more common in preclinical research it becomes critical for there to be a straightforward, easily replicable, accurate methodology for the non-biomedical optics community to apply fluence compensation to photoacoustic images. This methodology needs to be user-friendly and not require an extensive knowledge of the field of laser-tissue interactions. We propose the following method of effortlessly performing fluence compensation on PA images with the representative case of a commercial preclinical imaging system, VisualSonics (VS), using the open-source Monte Carlo toolbox, MCXLAB[55, 78].

Our methodology incorporates a newly developed PHotoacoustic ANnotation TOolkit for MATLAB (PHANTOM) that assists in user segmentation of US images into 3D labeled tissue structures with assignable optical properties. In addition, we have replicated the optical fiber illumination of the VS system as a light source in MCXLAB that accounts for optical fiber size, focus depth, and frame of focus within a 3D scan. The scripts and helping functions are deployed as an open-source toolkit which can be easily accessed by other biomedical researchers for fluence compensation of PA images acquired with VS system. Below we demonstrate the effect in phantom studies involving a blood-filled tube with different levels of blood oxygenation placed below *ex vivo* tissue with varying thickness. The efficacy of the process was further evaluated using two distinct kinds of *in vivo* data, namely a subcutaneous tumor model and a placental model of ectopic calcification. We hope to encourage researchers in the other fields such as nanoparticle drug delivery systems, perfusion modeling, contrast agent generation, etc. where PAI has become a valuable tool, to be able to easily generate accurate PA data with PHANTOM.

## 2. Materials and Methods

### 2.1 Workflow

Applying fluence compensation to large datasets of photoacoustic images can be a tedious and time-consuming task. Herein we describe a workflow, shown in Fig. 1, for fluence compensating photoacoustic images generated from a commercial imaging system, Vevo LAZR-X (FujiFilm, VisualSonics) that can be adapted to other imaging systems under similar reflective mode illumination schemes. Once a photoacoustic image is acquired using this system, it can be exported from the VevoLab software to perform additional post-processing. For this workflow the export data format is ‘.xml’ files, which will export from VevoLab in the ‘.raw.xml’ file format. processRawData.m is one of two code files that the user will need to run from start to finish of this workflow. This script will prompt the user for the saved file and read in the post-beamformed ultrasound and photoacoustic data as matrices and co-register the two based on complementary spatial axes also imported from the ‘.xml’ files. The script will then call the custom-built graphical user interface (PHANTOM-GUI) to assist with image segmentation and MC simulation set-up. Video S1 displays the aforementioned workflow.

**Figure 1.**
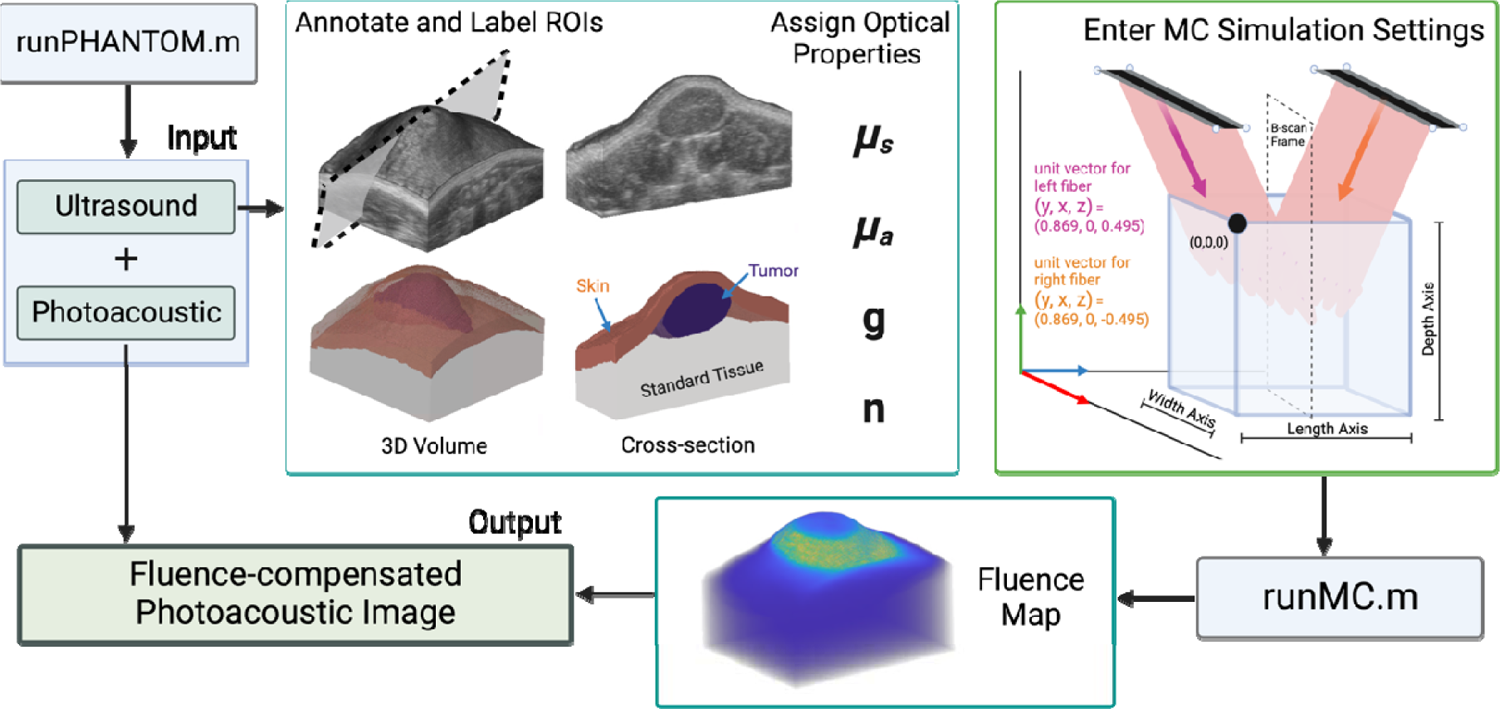
Illustration of the PHANTOM Workflow displaying the pipeline of how ultrasound data is used to create a labelled tissue volume with assigned optical properties. The beam geometry and MC simulation settings are set in relation to this volume and used to run MCX and generate a fluence map that is applied to the PA image for fluence compensation.

#### 2.1.1. Annotating Regions of Interest

The annotation panel of the PHANTOM-GUI offers extended functionality to users annotating 3D images. Scroll bar integration allows one to quickly view all frames within the 3D image stack and there are numerous annotation settings accessible to the user. For clarity, the term region of interest (ROI) will refer to the volumetric region of interest that is composed of 2D annotations. When starting a ROI, the user can select either a ‘2D’ or ‘3D’ option. These options both produce volumetric ROIs, but the 2D ROI will repeat a single annotation across all image frames, while a 3D ROI will have the user annotate each image frame individually. The 2D ROI is intended for temporal scans where the transducer remains stationary, while the 3D ROI is meant to be used for volumetric scans.

The user is also given the option of annotation shape. Users can select a rectangular, elliptical, or polygonal drawing tool to add flexibility. Most *in vivo* images will contain irregular shapes with smooth edges, so an interactive smoothing tool was also added. As volumetric scans are annotated frame by frame, an interpolation feature is included to speed up the annotation process. If the user has annotated a selection of frames of the imaging volume, a checkbox can be clicked to interpolate and generate annotations between user annotated frames. It is recommended for smooth interpolation that users annotate every 5-7 frames.

In the application of Monte Carlo simulations, it is often that multiple regions of interest need to be annotated as different tissue types can vary broadly in their optical properties. Due to this, the user can add as many ROIs as necessary, and assign each a label corresponding to the tissue type. As the user generates multiple ROIs, each will be tabulated according to the order in which they were drawn. The table of ROIs will also display the label name and give the option to interpolate between frames in the case of 3D ROIs. Users can easily toggle between each ROI by clicking on the corresponding row in the summary table or by utilizing the vertical scroll bar. Users can easily view the annotations on each frame as well as toggle between the different labeled ROIs. Users can export all of these different annotations and labels into a data structure for additional processing.

For the application of small animal imaging, an automatic segmentation algorithm was developed that enables the segmentation of the tissue region from the background region for all frames. The algorithm also allows segmentation of skin, given the user input of skin thickness in pixels. With the image dimensions being displayed on the image axes, this requires minimal effort from the user. The automatic segmentation of the general tissue structure allows the user to focus on annotations relevant to their work, such as defining tumor or organ regions. We hope that these features help to speed up the timely task of annotating large datasets.

#### 2.1.2. Monte Carlo Simulation Parameter Inputs

Once the user has completed all desired inputs, they can switch to the MC parameter tab to configure the simulation. To accommodate varying levels of experience in the field of biomedical optics, the PHANTOM-GUI offers the option of choosing the default optical properties or entering custom properties. The default optical properties of 14 generic tissue types were set as described by Jacques[20], which will be expanded upon in further sections. Several additional parameters requiring user input must also be set to configure the MCXLAB simulation including, number of photons to launch, wavelength of light, desired bin size, step size of the 3D scan, as well as the start time, end time, and time step for the simulation. The user may also select the fiber type used (blue or green) and the desired output provided by MCXLAB (fluence, flux, or energy). Once all fields have been filled, the user can export the MC configuration as a structure saved in the .mat file format. To accommodate the need for multiwavelength compensation, the user need simply change the wavelength field value and re-export.

#### 2.1.3. Running the MCXLAB Simulation

To run the simulations via MCXLAB, the user runs the script ‘runMC.m’, which will prompt for the aforementioned save file and automatically run simulations for each image frame. The function will return the normalized 3D fluence, flux, or energy map, based on the output type selected, along with the 3D fluence compensated PA image.

### 2.2 Image Processing Algorithms

#### 2.2.1 Tissue Segmentation

The algorithm used to segment the tissue region from background region takes a 2D or 3D ultrasound image as input. The image is rescaled to range from 0-1 and a median filter is applied. Non-linear mapping of the image in the range 0.15-1 to the range 0-1 is applied using a gamma correction factor of 0.65. The image is then segmented into two regions using Simple Linear Iterative Clustering[79] and binarized. Each frame of the binary image is then blurred via 2D convolution with a kernel of 15×15 pixels and smoothed using simple thresholding. Sobel edge detection[80] is then used to determine the tissue boundary, and all pixels on and below this boundary are labeled as 1 (foreground) or 0 (background). This output binary image represents the tissue region and background. This technique is the basis for the automatic tissue, skin, and background segmentation options available to the user of PHANTOM.

#### 2.2.2 ROI Interpolation Algorithm

In the context of ROI interpolation, R_k_ and R_k+1_ are two user annotated frames and F is the number of blank frames between them. Assume that R_k_ and R_k+1_ are two successive slices of a region (R) contained in a digital binary volume (V_b_). The goal is to interpolate F number of slices R_k1_, R_k2_, …, R_kF_, between the slices of R_k_ and R_k+1_. Estimations of R are contained within these interpolated slices[81]. The Euclidean distance transform of the perimeter of R_k_ and R_k+1_ is calculated and each pixel in this transform is set to −1 outside the annotation boundary and to 1 inside the annotation boundary. Three-dimensional linear interpolation is then applied to R using the built in MATLAB function ‘interp3’ to create a denser region R’ with F+2 frames where the first slice is R_k_, the last slice is R_k+1_, and the middle slices are slices R_k1_, R_k2_, …, R_kF_. The final 3D binary image (B) is defined as: B(x,y,z) = 1 where R’(x,y,z) = -|R’(x,y,z)|. This last condition ensures that the returned matrix is binary with no negative values.

### 2.3 Default Optical Properties

#### 2.3.1. Optical Absorption Coefficient

The absorption coefficient of a generic tissue type can be described by the relative contributions of the dominant chromophores contained within the tissue as described by Jacques[20]. The tissue parameters of average blood volume fraction (B), blood oxygen saturation (S), tissue water content (W), melanin volume fraction (M), and tissue fat content (F) can be used to specify the expected absorption coefficient of any generic tissue type when given the known absorption coefficients of hemoglobin (µ_a.oxy_, µ_a.deoxy_), water (µ_a.water_), melanin (µ_a.melanosome_), and fat (µ_a.fat_), as shown in EQ 1.

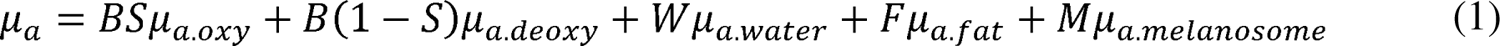

The values of B, S, W, F, and M used to calculate the absorption spectrum for the generic tissue types provided are tabulated in Table S1.

#### 2.3.2. Reduced Scattering Coefficient

As described by Jacques[20], the relationship between wavelength and reduced scattering coefficient (µ_s_’) can be described by EQ 2, where α’ represents the reduced scattering coefficient of the tissue at a 500 nm wavelength which is used to scale the wavelength dependent terms, λ is the wavelength (in nm), f_Ray_ is the Rayleigh scattering fraction and b_Mie_ represents the decreasing power for the Mie scattering component.

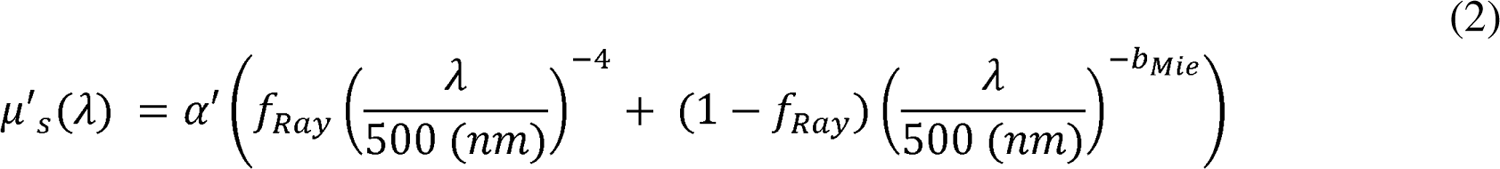

The values of α’, f_Ray_, and b_Mie_ used to calculate the reduced scattering coefficient (s) for the generic tissue types used in this work are tabulated in Table S2.

### 2.4 Modeling Light Distribution and Beam Geometry

The bifurcated light source was modeled using MCXLAB based on the physical dimensions of the optical fibers. Measurements of the fiber’s dimensions and position relative to the transducer were taken using digital calipers. The bifurcated fibers were modelled to be 10 mm apart in the z-direction at an angle of 20° from normal with the fiber focus at a depth of 8-12 mm from the transducer. The length of the fibers in the x-direction was modelled based on the various fiber types available for the Vevo LAZR-X system with blue, and green fibers being set at 24, and 14 mm respectively. A schematic visually displaying the blue optical fiber beam geometry is included in Fig. S1. The fiber width in the z-direction was measured using our own system with digital calipers and set to be 1.00 for the blue and 1.25 mm for green fibers.

Fibers were modeled as two planar sources defined by a position and direction. For clarity, when referring to the imaging volume and its surroundings, the x-direction will be the column-wise direction representing the width axis, the y-direction will be the row-wise direction representing the depth axis, and the z-direction will be the frame-wise direction, representing the scan direction axis. The coordinate grid encapsulating the imaging volume has the origin set as the uppermost left forward position. As the fibers are positioned above the imaging volume, the y-position of the fiber source will be negative.

The position of each planar source is defined relative to the imaging volume and is specified using corner points, while the direction is defined using unit vectors. As each US-PA image has an associated spatial width and depth relative to the transducer, the width and depth axes of the images can be used to determine the light source position. It is also important to note that for volumetric scans, the transducer position will change relative to the volume for each frame of acquisition. The pair of unit vectors used to describe source directions were (0.869, 0, 0.495) for the fiber shown on the left and (0.869, 0, −0.495) for the fiber shown on the right. A detailed schematic of the light illumination geometry and source coordinate positions in relation to the imaging volume can be found in Fig. S2. Prior to performing a simulation, the voxel size of all 3 dimensions needs to be equal for the accurate modeling of photons through each voxel. Three-dimensional interpolation was used to scale each dimension of the image so that each voxel has a length of the user-input bin size in each direction.

### 2.5 Phantom Experiments

#### 2.5.1. Phantom Preparation

To assess the accuracy of the optical fiber modelling used for MC simulations, a simple phantom experiment was conducted. This phantom consisted of blood in a tube with a section of chicken breast tissue placed on top. The chicken tissue had a gradient thickness ranging from 0 to ∼3 mm. A simple phantom mold was created using a 11 by 7.5 by 3 cm plastic box. Two 2 mm holes were melted into the box utilizing a soldering tool. The holes were positioned at a height of 2 cm from the bottom of the box and width of 3.25 cm along the short side of the box on each side. A clear 2 mm diameter (ID: 1.4 mm, OD: 1.9 mm) polyethylene tube (PE200, Intramedic™) was run through each hole and held taut while any excess space was filled using hot glue.

The phantom was prepared from porcine skin gelatin powder (G2500, Sigma–Aldrich) at a concentration of 8% (w/v). The gelatin powder was mixed with water at 50°C and stirred with heating until the gelatin completely dissolved. Once the temperature reached 60°C, the heat was removed and stirring continued until the gelatin reached a temperature of 40°C. The gelatin was poured into the phantom mold until covering half the surface area of the tube and air bubbles were removed using an aspirator. The mold was placed in 4°C fridge while the remaining gelatin solution cooled to 30°C with constant stirring. It is important to note that the temperature of 30°C was the maximum temperature allowable to prevent cooking of the chicken breast within the gelatin. Once 15 minutes had passed, the chicken breast was placed parallel to the tube and the remaining gelatin was poured into the mold leaving a height of 1 mm above the topmost section of chicken breast. An aspirator was used to remove any visible air bubbles on the surface and underneath the chicken breast. The phantom was left to solidify in 4°C fridge for 3 hours prior to imaging. Blood was prepared using powdered bovine hemoglobin (H2500, Sigma-Aldrich) dissolved in 1X phosphate buffered saline at a concentration of 2.5 mM. The blood solution was determined to be fully oxygenated after preparation using an oxygen monitor (Oxylite^TM^ Pro, Oxford Optronix).

#### 2.5.2. Phantom Imaging

All photoacoustic data was acquired using the Vevo LAZR-X small animal imaging system at the wavelengths of 750 and 850 nm using the MX250S transducer with a 6 dB bandwidth of 15-30 MHz and a center transmit frequency of 21 MHz The PA gain was set to 40 dB and the US gain was set to 10 dB. The persistence setting was on high, meaning each frame is a composite of ten averages. The step size of each 3D scan was 0.152 mm. All scans were acquired in triplicate on the same date and the scan started where there was no tissue inclusion above the blood tube and continued throughout the tissue inclusion, with the tissue thickness increasing with each subsequent frame. A schematic of the phantom imaging set-up is shown in Fig. 2.

**Figure 2.**
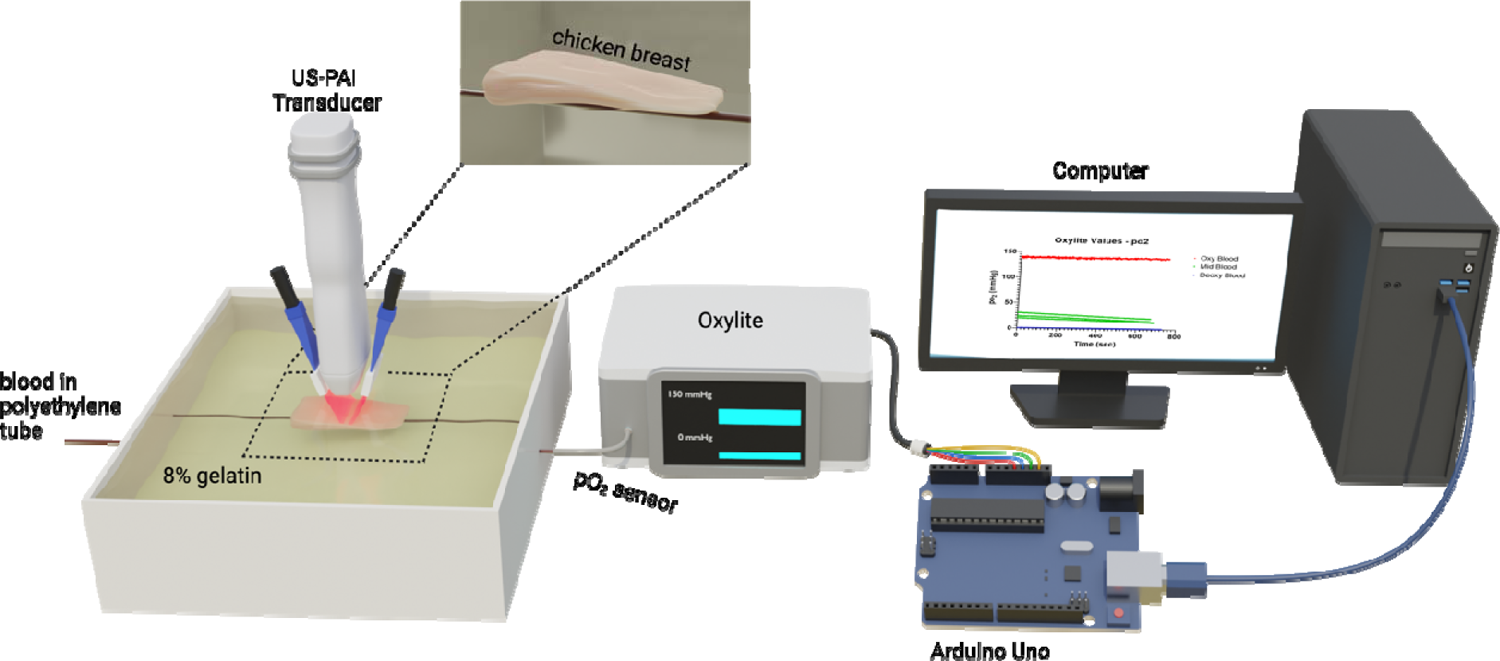
Schematic displaying the phantom imaging set-up. The oxygenation state of the blood-filled tube was analyzed with an oxygen monitor connected to a microcontroller board to directly send analog output into MATLAB in real-time. Data acquisition for blood pO_2_ occurred simultaneously with all photoacoustic scans.

Briefly, the oxygenated blood scans were performed first to ensure that no sodium dithionite was present in the tube. The tube was flushed with water between each scan. The deoxygenated blood scans were performed next, and blood was taken from the stock solution and deoxygenated with 2.5 mg/mL sodium dithionite. For the semi-oxygenated scans, 1.7 mg/mL sodium dithionite was added to blood taken from the stock solution. The blood was monitored with the oxygen monitor until reaching ∼60% StO_2_ when the first scan was begun. The three scans were taken as the blood was continuously undergoing deoxygenation. The oxygen monitor was connected to a microcontroller board (Arduino Uno, Arduino) to directly send analog output into MATLAB (MathWorks) in real-time. Data acquisition for blood pO_2_ occurred simultaneously with all photoacoustic scans. The raw pO_2_ values used to calculate the ground truth StO_2_ in the phantom experiments are provided in Fig. S3. As the blood used in the phantom experiments for each oxygenation state came from the same stock solution, this would result in the hemoglobin concentration being constant throughout the tube regardless of position or oxygenation state.

#### 2.5.3. Fluence Compensating Phantom Data

Each US image from the phantom scans was segmented and labeled utilizing PHANTOM. The chicken breast inclusion was assigned custom properties for µ_a_ and µ_s_’. The absorption coefficient for the chicken breast was calculated for both wavelengths using EQ 1, given then blood, water, and melanin content of muscle from Jacques with blood oxygen saturation set to 0%. The reduced scattering coefficient was calculated using EQ 2 for both wavelengths with the value of alpha taken from Marquez[82] and the values of f_Ray_ and b_Mie_ for muscle from Jacques. The optical properties of the background (gelatin) were assigned to equal those of water. For the deoxygenated and oxygenated blood scans, the spectra of Hb and HbO_2_ as compiled by Prahl[83] were used for the absorption and scattering coefficients of blood at both wavelengths. The optical properties of blood used for the semi-oxygenated scans were varied at each scanning frame based on the StO_2_ value calculated from the real-time pO_2_ measurements of the blood in the phantom. The optical properties for each label are given in Table 1.

**Table 1.**
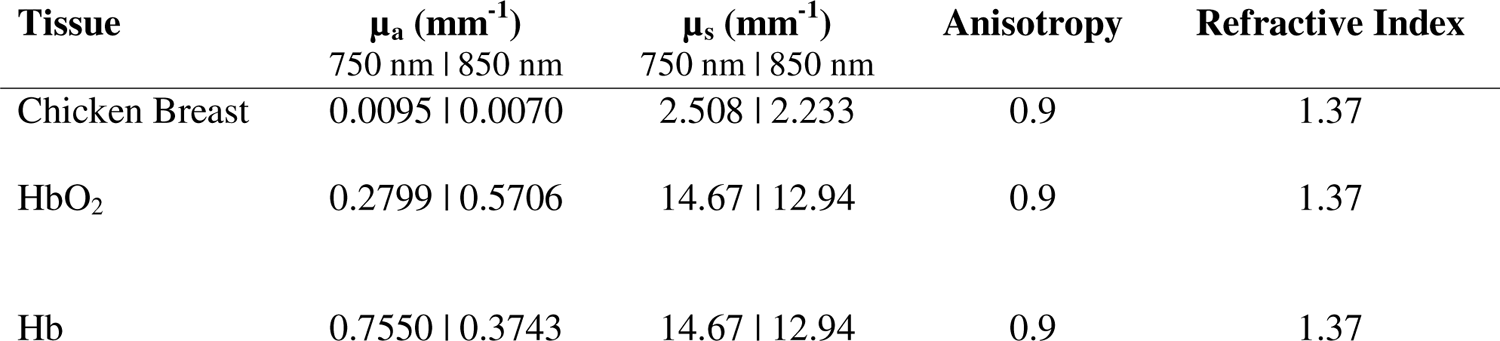
Optical properties used to fluence compensate phantom data.

| Tissue | $\mu_a$ (mm<sup>-1</sup>)<br>750 nm \| 850 nm | $\mu_s$ (mm<sup>-1</sup>)<br>750 nm \| 850 nm | Anisotropy | Refractive Index |
| --- | --- | --- | --- | --- |
| Chicken Breast | 0.0095 \| 0.0070 | 2.508 \| 2.233 | 0.9 | 1.37 |
| HbO<sub>2</sub> | 0.2799 \| 0.5706 | 14.67 \| 12.94 | 0.9 | 1.37 |
| Hb | 0.7550 \| 0.3743 | 14.67 \| 12.94 | 0.9 | 1.37 |

### 2.6 StO_2_ Calculations

#### 2.6.1. Spectral Unmixing of PA Images

As the two wavelengths used to generate the photoacoustic images in this study are 1 and 2, the molar concentrations of Hb and HbO_2_, can be linearly unmixed by solving a non-negative linear least-squares optimization of EQ 3 [25], where PA represents the photoacoustic image. This was calculated using the function ‘lsqnonneg’ from the MATLAB Optimization Toolbox.

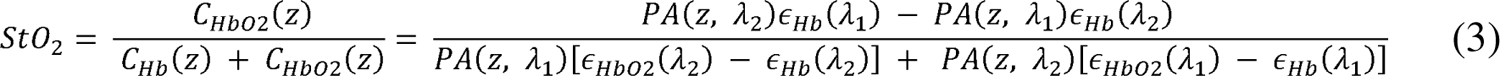

#### 2.6.2. Thresholding of StO_2_ from Normalized HbT Signal

All HbT data presented in this manuscript was normalized based on the maximum HbT signal throughout the imaging volume. This allowed meaningful comparisons between the HbT signal of non-compensated and fluence-compensated data. All StO_2_ values presented were thresholded to only include data from voxels where the normalized HbT was greater than or equal to 0.02. This condition ensures that StO_2_ values are being calculated from voxels where there is hemoglobin signal above the noise threshold.

#### 2.6.3. StO_2_ calculated from pO_2_ Data

The pO_2_ values gathered with the oxygen monitor during the PA phantom imaging were converted to blood StO_2_ values using the oxyhemoglobin dissociation curve relationship described by Severinghaus[84, 85], shown in EQ 4.

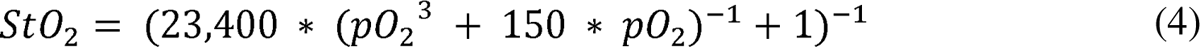

### 2.7 Validation in a murine subcutaneous tumor model

#### 2.7.1. Tumor Model

The Institutional Animal Care and Use Committee (IACUC) at Tufts University authorized all animal experiments conducted in this study. Male homozygous Foxn1nu nude mice (The Jackson Laboratory), aged 6-8 weeks, were and subcutaneously injected with 5 million AsPC-1 cells in 100 μL of Matrigel (50 μL of Matrigel + 50 μL of PBS) using a 1-ml tuberculin syringe with a 30-gauge hypodermic needle. Cells were obtained from American Type Culture Collection and maintained in RPMI (Roswell Park Memorial Institute) 1640 medium. AsPC-1 cells were passaged 1-2 times per week. Media were supplemented with 10% fetal bovine serum and 1% penicillin-streptomycin (100 U/ml). All cells were cultured in suspension and kept in an incubator in 5% CO_2_ at a temperature of 37C.

#### 2.7.2. Tumor Imaging

US and PA images were acquired using the Vevo LAZR-X using the MX250S linear array transducer. Gain was set to 22 dB for US, and 45 dB for PAI. Persistence was set to Max (20 averages per frame) and images were acquired with the Oxy-Hemo setting (750 and 850 nm laser illumination wavelengths). Mice sedated with isoflurane (2-3% induction, 1.5% maintenance) were put on a heating pad with ECG leads to monitor body temperature, heart rate, and respiration during the imaging session. Bubble-free ultrasound transmission gel (Aquasonic100 Ultrasonic Transmission Gel, Parker Laboratories, Inc.) was placed to the tumor to enhance acoustic transmission between the transducer and tumor during imaging. The 3D image of tumor StO_2_ values was obtained between 20 to 30 minutes, with the initial imaging frame recorded at the anterior end of the tumor and each subsequent frame 0.15 mm farther in the posterior direction.

#### 2.7.3. Fluence Compensation

The fluence compensation workflow was tested on PA images of a subcutaneous pancreatic tumor at 52 days post-implantation. This tumor had a volume of 379.5 mm^3^ with a depth of over 6 mm from the tissue surface at the deepest point. The tumor region was user annotated from the US image every 5 frames utilizing the interpolation feature of the PHANTOM-GUI. The background, skin, and tissue boundary were annotated using the automatic segmentation features with skin thickness set to 20 pixels, and manually edited by the user for specific frames with regions of skin with non-uniform thickness. The 4 labels used to assign optical properties to this volume were, water (background), skin, tumor, and standard tissue. The optical properties of water, skin, and standard tissue were assigned using the generic tissue types. The tumor optical properties were set using the default tissue properties for absorption, anisotropy, and refractive index, while the scattering coefficient was custom set based on EQ 2. Values of f_Ray_, and b_Mie_ were obtained from work by Sandell and Zhu[86], while alpha was obtained from work by Wilson et al.[87] The optical properties of all 3 tissue types are summarized in Table 2.

**Table 2.** Optical properties from Jacques[20] used to fluence compensate PA scan of subcutaneous tumor.

| Tissue | $\mu_a$ (mm<sup>-1</sup>) | $\mu_s$ (mm<sup>-1</sup>) | Anisotropy | Refractive Index |
| --- | --- | --- | --- | --- |
|  | 750 nm \| 850 nm | 750 nm \| 850 nm |  |  |
| Skin | 0.2504 \| 0.2935 | 25.22 \| 21.90 | 0.9 | 1.37 |
| Tumor | 0.0051 \| 0.0061 | 5.278 \| 4.578 | 0.9 | 1.37 |
| Standard Tissue | 0.0110 \| 0.0199 | 10.97 \| 9.418 | 0.9 | 1.37 |

**Table 3.** Optical properties used to fluence compensate PA scan of *in vivo* placenta.

| Tissue | $\mu_a$ (mm<sup>-1</sup>)<br>750 nm \| 850 nm | $\mu_s$ (mm<sup>-1</sup>)<br>750 nm \| 850 nm | Anisotropy | Refractive Index |
| --- | --- | --- | --- | --- |
| Skin | 0.2504 \| 0.2935 | 25.22 \| 21.90 | 0.9 | 1.37 |
| Placenta (liver) | 0.2083 \| 0.2729 | 8.486 \| 6.911 | 0.9 | 1.37 |
| Fetus (bone) | 0.0012 \| 0.0020 | 13.18 \| 12.63 | 0.9 | 1.37 |
| Standard Tissue | 0.0110 \| 0.0199 | 10.97 \| 9.418 | 0.9 | 1.37 |

### 2.8 Validation in a murine model of ectopic calcification

#### 2.8.1. Animal Model

All animal studies were conducted under the appropriate protocols approved by the IACUC at Tufts University. Wildtype (*Slc20a2*+/+) C57Bl/6J mice (Jackson Labs) and C57BL/6NTac-Slc20a2^tm1a(EUCOMM)Wtsi^/Ieg (*Slc20a2* +/−) mice (European Mouse Mutant Archive) were used to establish the colonies, mice, and placentas used throughout the experiments.

#### 2.8.2. Animal Imaging

A maternal model for (*Slc20a2*+/-) was analyzed. At least 12 hours prior to imaging the animals’ abdomen, fur was clipped down to make depilation easier and avoid chemical burns from depilation cream. The animals were imaged under anesthesia on gestational day 18.5 under 1-1.5% isoflurane. Image collection was performed using the Vevo LAZR-X ultrasound and photoacoustic system. *In vivo* US-PA images were collected at 750 and 850 nm wavelengths and a 21 MHz transducer (MX250S). The PA and US gain were set at 45 and 22 dB, respectively. US was first utilized to approximate the number and location of visible embryos and placentas followed by a 3D US-PA scan of 1 superficial placenta.

#### 2.8.3. Fluence Compensation

The fluence compensation workflow was also tested on a 3D PA of a gestational day 18.5 placenta. This placenta had a volume of 114.4 mm3 and maximum depth of ∼8 mm. The placenta and embryo region were user annotated using PHANTOM-GUI, while the skin, background, and tissue region were annotated using the automatic segmentation feature of the PHANTOM-GUI. The 5 labels used to assign optical properties to this volume were, water (background), skin, placenta, embryo, and standard tissue. The optical properties of µ_s,_ anisotropy, and refractive index for all labels were defined using the generic optical properties provided in PHANTOM, where the placenta was approximated as liver, and the embryo was approximated as bone. The value of µ_a_ for water, skin, bone, and standard tissue were set using the default optical properties, while the value of µ_a_ for placenta was custom set using EQ 2 with values for blood volume fraction, water content, fat content, and melanin content taken from Badachhape[88], Lee[89], Pratt[90], and Yi[91] respectively.

### 2.9 Histology

Post imaging, pimonidazole HCl (Hypoxyprobe Inc) was administered at a concentration of 60 mg/kg via tail vein injection and mice were sacrificed one hour later. Tumors were surgically excised without skin and then embedded in optimal cutting temperature (OCT) compound (Tissue-Tek) in the same alignment as the US-PAI cross-sectional images. The cryo-sections of tumor tissue, with a thickness of 10 microns, were meticulously prepared utilizing a cryotome and subsequently affixed onto glass microscope slides. Hematoxylin and eosin (H&E) and immunofluorescence staining procedures were employed for histological examination as previously described[28].

The cryo-sections were initially subjected to fixation by immersion in a pre-cooled mixture of acetone and methanol at a 1:1 ratio for a duration of 10 minutes while maintained at an ice-cold temperature. Subsequently, the sections were air-dried for 30 minutes and then subjected to three sequential 5-minute washes with 1x phosphate-buffered saline (PBS). Following this, a blocking solution with a 1x concentration (BSA, Thermo Scientific™) was applied to the sections, allowing incubation period of 1 hour at room temperature.

For the immunostaining of microvasculature and hypoxic regions within the tumor sections, primary antibodies, namely Mouse CD31/PECAM-1 Affinity Purified Polyclonal Ab (R&D Systems Inc) and FITC-Mab (Hydroxyprobe Inc), were incubated with the tissue sections at an approximate concentration of 10 μg/mL. The incubation of these antibodies was carried out overnight at 4 °C. On the subsequent day Donkey Anti-Goat IgG NL493 Affinity Purified PAb (R&D Systems Inc) secondary antibody was introduced following washing with 1x PBS. This secondary antibody was allowed to incubate at room temperature for 2 hours. Slowfade gold antifade mountant containing 4’,6-diamidino-2-phenylindole (DAPI, Invitrogen) was used to rinse and seal the coverslips. The acquired slides were imaged at 4x magnification utilizing an EVOS fluorescence imaging system.

### 2.10 Statistical Analysis

GraphPad Prism (La Jolla, CA) was used for all statistical analysis described in this manuscript. In all statistical tests used, a p-value less than or equal to 0.05 was deemed statistically significant.

## 3. Results and Discussion

### 3.1 Depth-dependent fluence compensation increases PA signal and accuracy of measured StO_2_

HbT signal strength for non-fluence compensated data decreases as a function of the tissue thickness above the tube for all 3 blood oxygen states, as expected and demonstrated in Fig. 3A (left columns for each case). Even at the tube locations with no chicken breast inclusion in the frame of focus (Fig. 3A top row), light attenuation is still present within the tube itself as seen by the higher amplitude signal at the upper portion of the tube than the lower part. Hemoglobin is a predominant absorber in the NIR range, resulting in the rapid attenuation of light and corresponding decrease in signal strength. As the thickness of the tissue increases, fluence reaching the tube is decreasing and therefore the signal from the upper portion of the tube diminishes, while the signal from the lower portion of the tube almost disappears entirely. Fluence compensation of this data decreases the attenuation seen as a function of tissue thickness as well as provides stronger signal at the lower portion of the tube (Fig. 3A, right column for each case). The decrease in HbT signal strength of the non-compensated data is quantitatively displayed (Fig. 3B). As the tube was filled with a homogeneous solution of *ex vivo* blood it can be assumed that the hemoglobin content was equal throughout the tube and the decrease in signal strength is the direct result of light attenuation. As light attenuation is known to be exponential according to the Beer-Lambert Law [92], the exponential fit (Fig. 3B, solid black line) was calculated for all 3 non-compensated curves and showed good agreement with the data (R^2^ = 0.96, 0.91, 0.96 for oxygenated, semi-oxygenated, and deoxygenated blood). Fluence compensating the data (Fig. 3B, blue dots) flattened this curve, showing a consistent measurement of HbT as tissue thickness increased.

**Figure 3.**
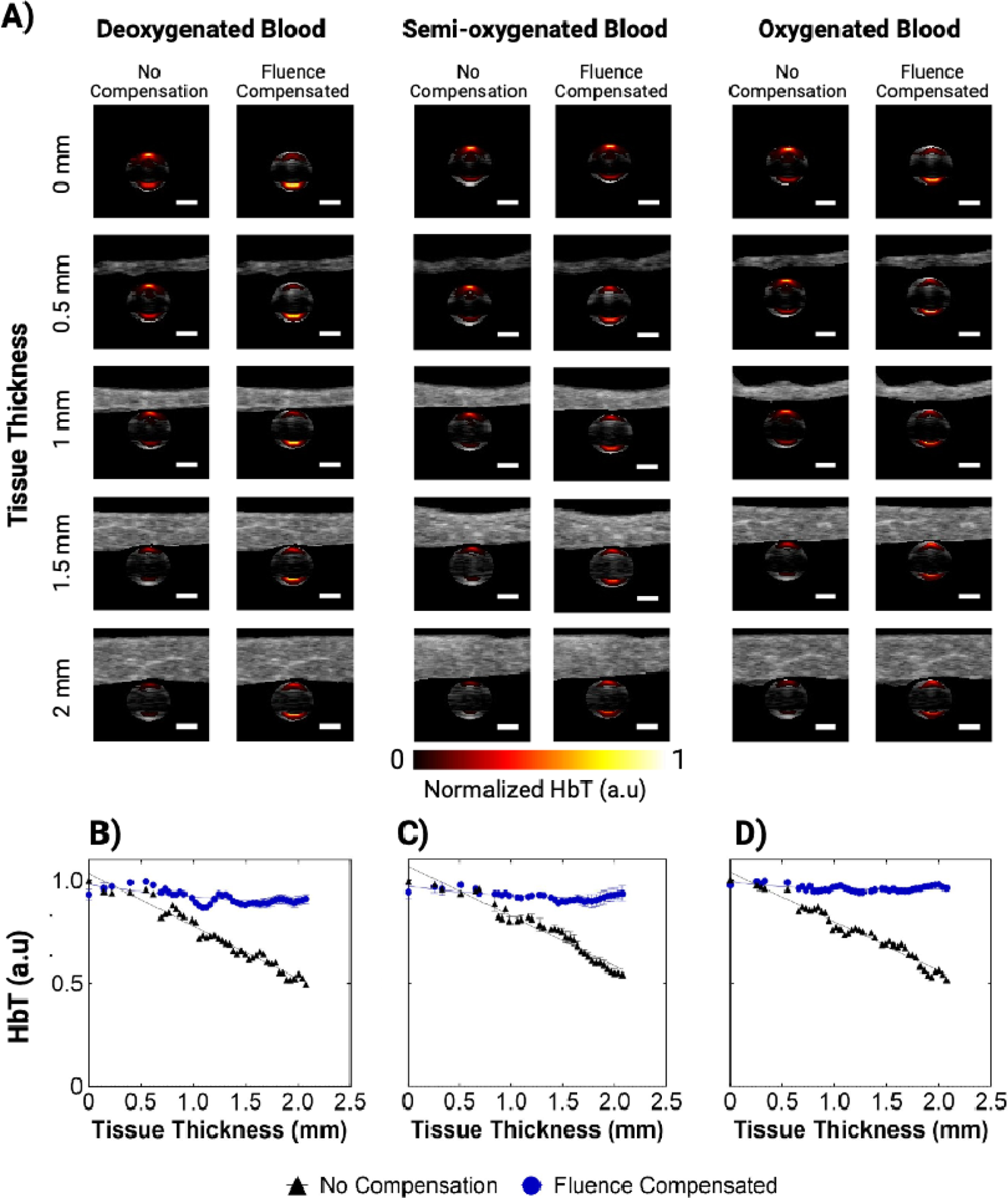
A) 2D rendering of the phantom showing US of the chicken breast inclusion and US-PA of the blood tube where the PA image shows the HbT normalized from the maximum value in the 3D volume. The HbT signal for deoxygenated (left), semi-oxygenated (middle), and oxygenated blood (right) are shown at 5 different frames for the compensated and non-compensated images, each corresponding to a location in the phantom with varying heights of the chicken breast inclusion. The scale bar shown represents 1 mm. B-D) Plots of HbT signal

In addition to HbT, StO_2_ is an invaluable prognostic marker gathered from multi-wavelength photoacoustic imaging for diagnosis and monitoring of diseases[27, 64, 70, 93].The StO_2_ data of the blood inside the phantom is visualized in 2D and 3D in Fig. 4A and Fig. 5 respectively. As seen in the 2D and 3D non-compensated images, there is a visible difference between the StO_2_ values measured at the bottom versus top region of the tube. This phenomenon can be attributed to the difference in light attenuation due to the optical properties of hemoglobin between 750 and 850 nm wavelengths. The StO_2_ data from the phantom experiment is quantitatively displayed (Fig. 4B) and shows that fluence compensating the phantom data improved the accuracy of the blood StO_2_ images when compared to the ground truth pO_2_ readings. Prior to fluence compensation, the StO_2_ values measured for deoxygenated blood had a mean value of 17.28% (σ=2.52, 95% CI = 16.51-18.04%) minimum of 13.84% and maximum of 21.83% with a range of 7.99%. These values are relatively higher than the StO_2_ values calculated from the fluence-compensated data which had a mean value of 3.77% (σ=0.81, 95% CI = 3.52-4.01%) with a minimum of 2.80%, maximum of 5.11%, and range of 2.31%. An unpaired two-tailed t-test with Welch’s correction for unequal standard deviation was conducted comparing the experimental error of the two groups. The t-test showed that there was a significant difference between the error (p-value < 0.0001, t = 30.15) with the fluence-compensated data showing less error (ΔxLJ= −19.33 ± 0.641) when compared to the ground truth. The experimental error for the deoxygenated blood condition is relatively very high due to the very small (close to zero) ground truth data.

**Figure 4.**
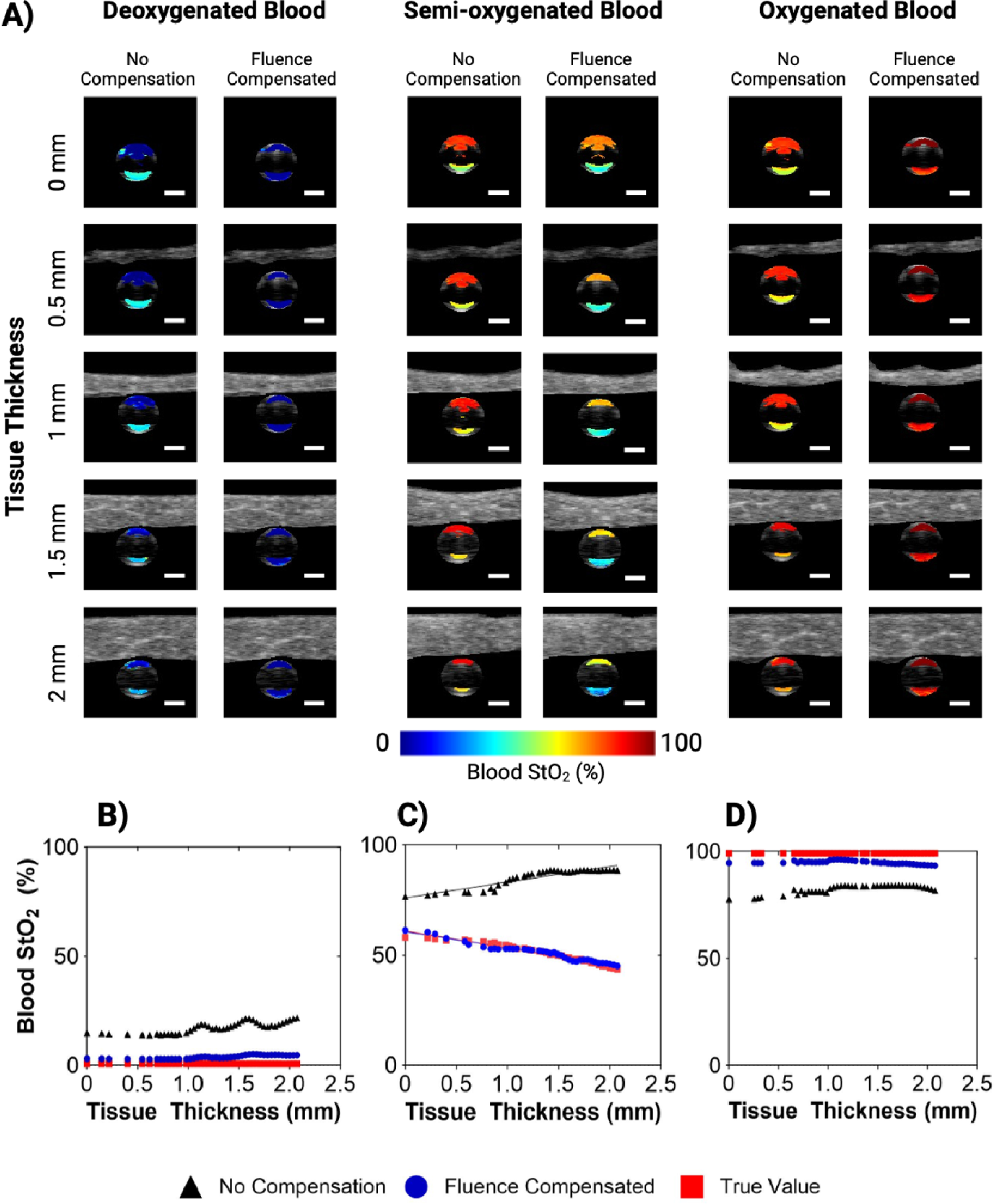
A) 2D rendering of the phantom showing US of the chicken breast inclusion and US-PA of the blood tube where the PA image shows the blood StO_2_. The StO_2_ values for deoxygenated (left), semi-oxygenated (middle), and oxygenated blood (right) are shown at 5 different frames for the compensated and non-compensated images, each corresponding to a location in the phantom with varying heights of the chicken breast inclusion. The scale bar shown represents 1 mm. B-D) Plots of StO_2_ against the thickness of the chicken breast inclusion from the phantom experiment for the deoxygenated (B), semi-oxygenated (C), and oxygenated (D) blood conditions. The fluence compensated data (FC) is shown in black, the non-compensated data (NC) is shown in blue, while the ground truth values (GT) are plotted in red. The linear fit for the NC, FC, and GT data are also plotted for the semi-oxygenated blood. R^2^ = 0.860, 0.960, 0.950 for NC, FC, and GT respectively.

**Figure 5.**
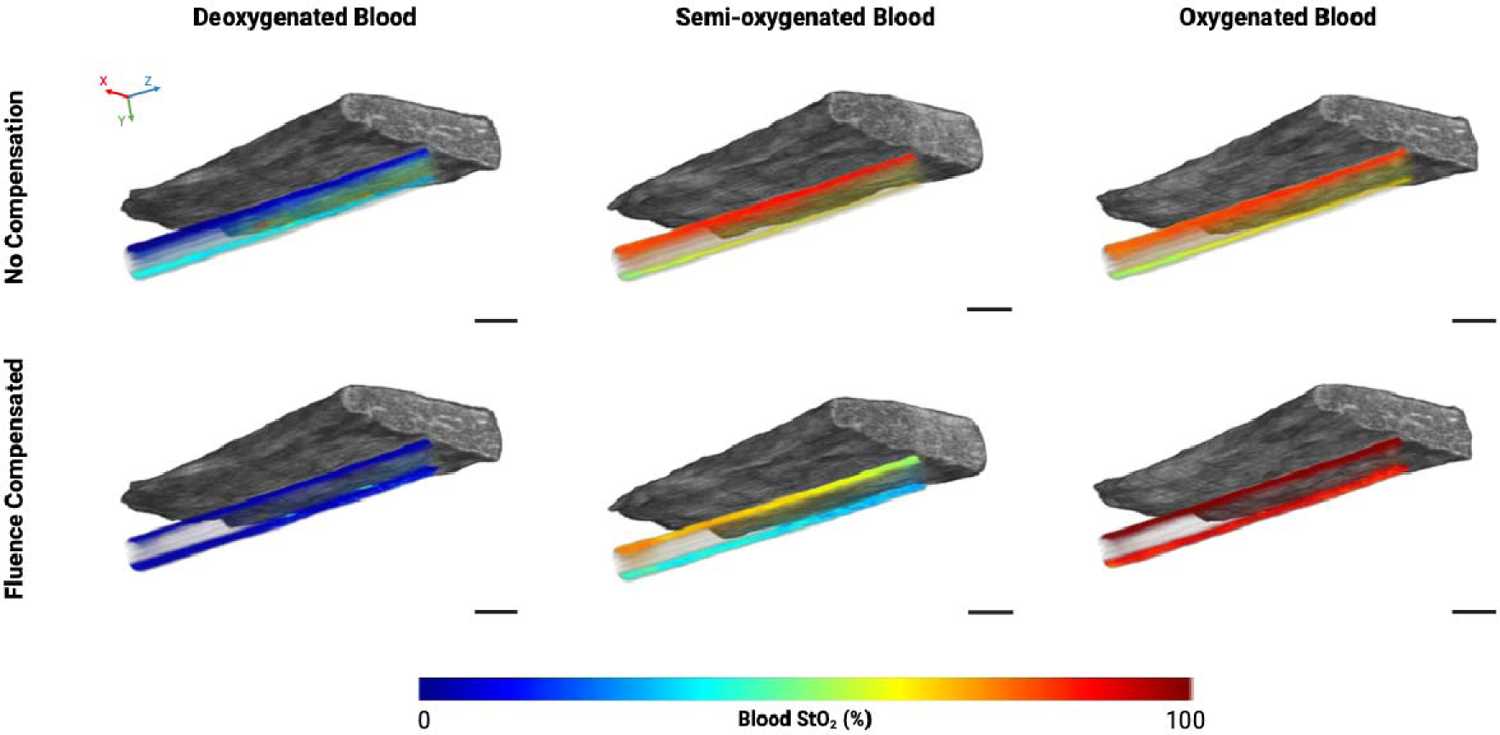
3D rendering of the phantom showing US of the chicken breast inclusion and PA of the blood tube where the PA image shows the blood StO_2_. The StO_2_ values for deoxygenated (left), semi-oxygenated (middle), and oxygenated blood (right) are shown for the compensated and non-compensated images. X, Y, and Z-direction are shown with red, green, and blue arrows respectively, where the Z-direction is the transducer scanning direction. The scale bar shown represents 2 mm.

The estimation of StO_2_ data for oxygenated blood was also improved with fluence compensation. A two-tailed t-test with Welch’s correction conducted to compare the experimental error of the non-compensated versus fluence compensated data found that there was a significant difference between the error of the two groups (p-value < 0.0001, t = 38.76) with the fluence-compensated data showing lower overall experimental error (ΔxLJ= −0.125 ± 0.003). The non-compensated StO_2_ values for oxygenated blood had a mean of 82.37% (σ=1.93, 95% CI = 81.78-82.97%), a minimum of 77.65%, maximum of 84.04%, and range of 6.39%. The compensated StO_2_ values had a mean of 94.76% (σ=0.80, 95% CI = 94.51-95.00%), minimum of 93.29%, maximum of 96.18%, and range of 2.88%.

The scans for each blood oxygenation state were conducted in triplicate, however the StO_2_ data for the three scans performed on the semi-oxygenated blood phantom were statistically analyzed separately as the StO_2_ values were fluctuating during each scan due to the ongoing deoxygenation reaction between the blood solution and sodium dithionite. The data for one representative scan is shown in Fig. 4B, while the plots for the remaining two scans can be found in Fig. S4. The addition of sodium dithionite at a concentration of 2.5 mg/mL was found to hold blood at 0% StO_2_ for almost one hour[94], allowing the deoxygenated blood scans to have a constant StO_2_. For semi-oxygenated blood however, the blood StO_2_ could not be held constant at one value, and scans were performed whilst the blood was still undergoing the deoxygenating reaction and actively decreasing in StO_2_ during each scan where the thickness of the chicken breast inclusion was increasing with each successive frame In fact, the StO_2_ of the blood in the tube was seen to slightly increase with increasing tissue thickness for the non-compensated StO_2_ values, which is the opposite trend to the ground truth StO_2_ decreasing throughout time (Fig. 4B-D). Once fluence compensated, the StO_2_ data reflects the ongoing chemical reaction within the tube, where the blood is actively undergoing deoxygenation.

ANCOVA analysis reveals that the slopes of the true value data and the fluence compensated data are not significantly different for all 3 scans (p-value = 0.428, 0.542, 0.126). The y-intercept was also found to have no significant difference between the fluence compensated and ground truth data for 2 of 3 scans taken (p-value = 0.752, 0.125, < 0.0001). The slopes of the non-compensated data and ground truth data and the non-compensated and fluence-compensated data were found to be statistically different from one another (p-value < 0.0001). While the blood was actively undergoing deoxygenation, the relative changes in StO_2_ were detectable using fluence compensated data, compared to non-compensated StO_2_ values. Analyzing the experimental error of the semi-oxygenated blood measurements for each scan with Welch’s t-test showed that there was a significant difference between the error of the two groups (p-value < 0.0001, t = 20.26), with the fluence compensated data showing lower average error (ΔxLJ ± SEM = −1.23 ± 0.061)

There are two noticeable trends displayed by this data. Firstly, we will discuss the effect of tissue thickness on StO_2_ (irrespective of blood oxygenation state). The non-compensated data displayed the trend of an increase in blood StO_2_ with increasing tissue thickness, which is clearly displayed by the 3D rendered images (Fig. 5). Theoretically, this trend complements itself well to the spectra of oxy- and deoxy-hemoglobin. For the case of homogenous blood StO_2_, longer wavelengths will penetrate deeper into the phantom. Therefore, with no chicken breast present above the tube, one can assume that after accounting for laser energy differences, equal amounts of 750 nm and 850 nm light are illuminating the top portion of the tube. When there are more scattering events above the tube, due to increasing tissue thickness, less overall light is hitting the top of the tube. It can also be assumed that with greater light attenuation due to the presence of the tissue, more 850 nm light is going to reach the tube compared to 750 nm. As oxygenated hemoglobin has a larger molar extinction coefficient than deoxygenated hemoglobin at 850 nm this will manifest as an observed increase in StO_2_ (Fig. 4B). In addition, the differences in light penetration for these two wavelengths are not expected to be drastically different, corresponding with the minor increase in StO_2_ displayed by this trend. It is important to note that blood StO_2_ values calculated from images of different wavelengths may not display this same trend.

The effect of light attenuation throughout the tube depth (irrespective of tissue thickness) is also noticeable in the non-compensated 2D and 3D StO_2_ images and differs based on blood oxygen state. For deoxygenated blood, the lower portion of the tube displays a higher average StO_2_ than the upper portion. When light is travelling through the depth of the tube, less absorption events occur at 850 nm due to the lower absorption coefficient of deoxyhemoglobin at 850 nm. Therefore, more 850 nm light will reach the lower area of the tube than 750 nm light, positively skewing the StO_2_ measurements in that area. When correcting for this unequal light attenuation with fluence compensation, the StO_2_ values are much more consistent throughout the entire depth of the tube as shown in Fig. 5. For the case of fully oxygenated blood, the result is opposite, and the lower portion of the tube displays StO_2_ values lower than the upper area. The oxyhemoglobin absorption curve is lower at 750 nm than it is at 850 nm, and the amount of 850 nm light reaching the bottom of the tube will be reduced due to light attenuation. This results in the calculated StO_2_ values at the bottom of the tube to be lower than the true value. The fluence compensation used in this work was able to correct for this to create a more uniform distribution of calculated StO_2_ for all depths within the tube.

### 3.2 Effects of Fluence Compensation on Subcutaneous Tumor Data

The normalized HbT and StO_2_ maps of the representative tumor are shown in 2D (Fig. 6A) and 3D (Fig. 6C) for the non-compensated and fluence compensated photoacoustic data. Qualitatively one can see that there is better signal strength at deeper areas of the tumor for the fluence compensated data than the non-compensated data in the 2D images, indicated by the white arrows. When viewing the 3D StO_2_ images (Fig. 6C, black arrows), one can visually see that there are areas of stronger signal strength throughout the whole volume in the fluence compensated image compared to the non-compensated image. The visual differences in the 3D renderings are quantitatively displayed in the volumetric distributions of HbT and StO_2_ for the compensated and non-compensated data (Fig. S5). While the shape of the HbT distribution remains visually similar for the fluence compensated and non-compensated images, a two-sided Wilcoxon rank sum test reveals that there is a statistically significant difference between the medians of the two distributions (p < 0.0001, z = 45.19). An unpaired two-sided t-test comparing the StO_2_ distributions of the compensated and non-compensated images indicates a statistically significant difference between the mean of the two distributions (p < 0.0001, t = 208.3). The shape of the StO_2_ distribution varies drastically between the un-compensated and fluence compensated data. There is a high concentration of zeros in both StO_2_ distributions that can be attributed to the non-negativity constraint applied to spectrally unmix the data, which will inherently shrink values towards zero. The uncompensated StO_2_ distribution is relatively flat with a small peak at around 85% StO_2_, while the compensated StO_2_ distribution has a more normal shape and displays a distinct peak at 47% StO_2_. The differences in these two distributions clearly indicate that a significant amount of data on the intertumoral heterogeneity is lost when StO_2_ measurements are calculated from non-compensated PA images.

**Figure 6.**
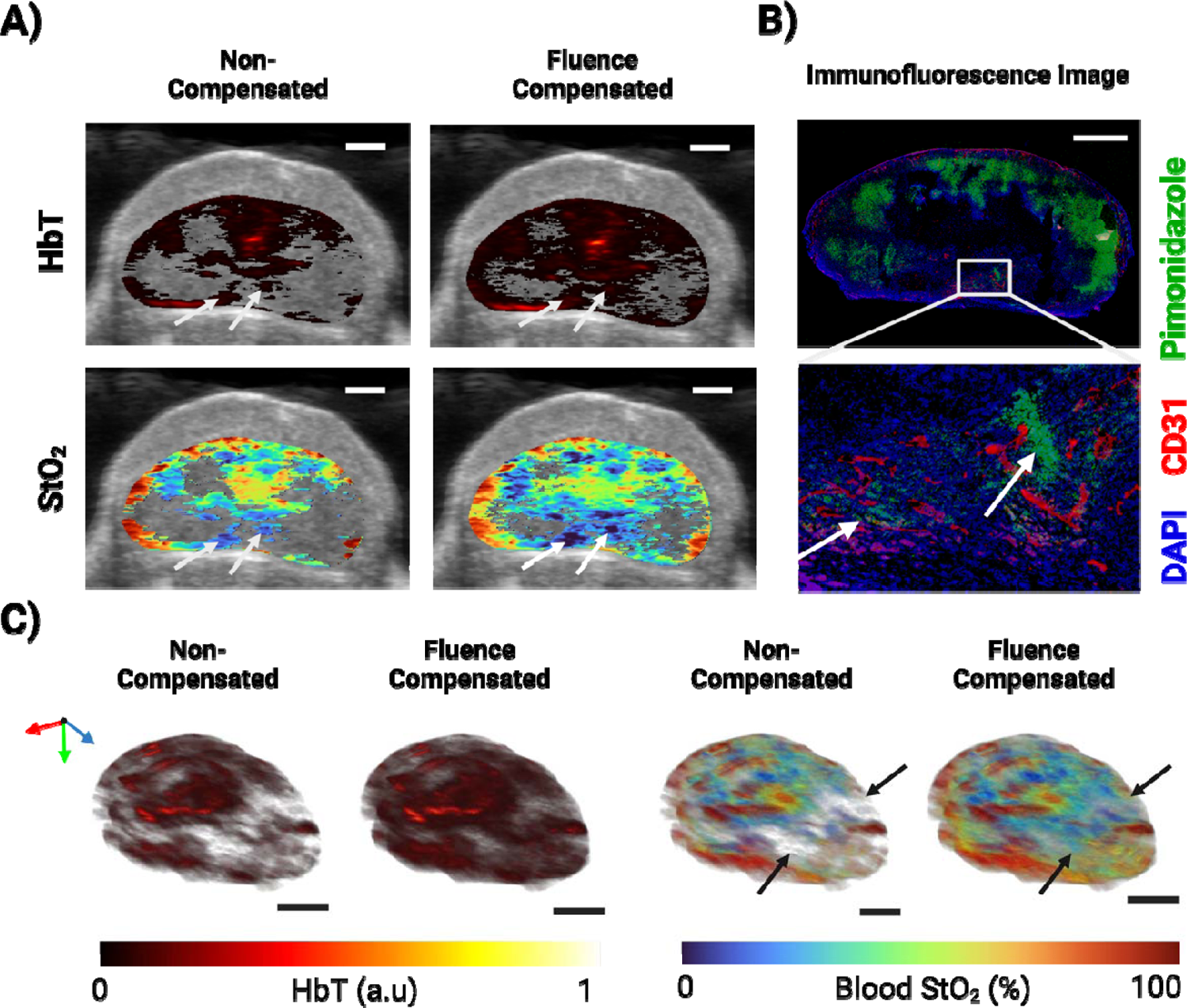
A) 2D renderings of tumor HbT (top) and StO_2_ (bottom) for non-compensated (left) and fluence compensated (right) data. B) Immunofluorescence stain for cell nuclei in blue, blood vessels in red, and hypoxia in green. stain of the tumor cross-section. C) 3D renderings of tumor HbT (top) and StO_2_ (bottom) for non-compensated (left) and fluence compensated (right) data. X, Y, and Z-direction are shown with red, green, and blue arrows respectively, where the Z-direction is the transducer scanning direction. The scale bar shown for all images represents 2 mm.

Comparing the 2D PA images with immunofluorescence (IF) image of the same tumor cross-section reveals that there is a highly vascularized hypoxic region at the bottom of the tumor (Fig. 6B - boxed in white). The corresponding region in the PA image has lower StO_2_ and higher HbT values in the compensated images compared to the non-compensated images. This is especially noticeable in the hypoxic areas with vascular signal as pointed out by the white arrows (Fig. 6A-B respectively). Fluence compensation gives results that show lower StO_2_ in regions of hypoxia. Of particular note, the compensated StO_2_ images show vascularized hypoxic areas with <10% StO_2_ that correspond well with the IF stain. This distinction is relevant as binding of the hypoxia marker, pimonidazole, increases drastically below 10 mmHg [95], which corresponds to an StO_2_ value of 9.7%.

The importance of accurate PA-based HbT and StO_2_ data in pre-clinical cancer research cannot be overstated. Fluence compensation greatly increases the reliability of StO_2_ measurements used to identify hypoxic regions, which has the potential to aid cancer diagnosis, treatment planning, and therapy monitoring. Pathological hypoxia exerts influence on individual cancer cells as well as the whole tumor microenvironment and regulates the key processes of vascularization, immune-resistance, cell death, and cell proliferation[96–98]. Hypoxia is a prominent characteristic seen in solid tumors and is widely acknowledged as a significant contributor to worse prognostic outcomes in individuals with cancer[99, 100]. By assessing tumor vasculature and the changes in oxygen levels, it is possible to infer if the tumor is undergoing hypoxia as a response to therapeutic intervention, or that it is enhancing its glycolytic metabolism and angiogenesis to facilitate proliferation and metastasis[101]. In order for physicians to select the most effective treatment and make educated choices throughout therapeutic intervention and follow-up it is essential to understand these aspects of tumoral activity.

Fluence compensation is particularly important in longitudinal studies that monitor changes in tumor hemodynamics over periods of weeks to months. In such studies, untreated or recurring tumors can become relatively large where the effects of light attenuation need to be accounted for. Areas of hypoxia or low vascular density could be incorrectly identified in large tumors due a diminished signal strength at depth if photoacoustic signals detected are not properly fluence compensated. In addition, this toolkit allows annotation of an unlimited number of regions. Fiducial markers can be annotated across images in specific tumor sub-regions, allowing for researchers to track regional changes in tumoral heterogeneity throughout treatment and monitoring. Distinct areas in or above the skin, such as sutures or ulcers/hemorrhages could also be annotated and to increase the intricacy of the label-based volume, given that the optical properties of these regions are known.

### 3.3 Effect of Fluence Compensation on Placental Model of Ectopic Calcification

Compared to the subcutaneous pancreatic tumor, the *in vivo* placenta displays high levels of vascularization as the placenta is an essential organ that facilitates the transfer of nutrients and oxygen between the maternal and fetal compartments. The placenta imaged was located lateral to the embryo just below the skin surface, as shown in Fig. 7A. Within the mouse placenta, there are three distinct regions as depicted in Fig. 7B: decidua (maternal tissue), junctional zone (fetal tissue), and labyrinth (fetal tissue) [102]. Visually, the maternal spiral arteries would be centralized into one or several channels that then branch through the junctional zone to create a complex tangle of maternal blood vessels in the labyrinth. Fetal capillaries are also branched and tangled within labyrinth to allow gas and nutrient exchange. This creates a mesh of blood vessels. Fig. 7C-D displays the 2D and 3D normalized HbT and StO_2_ maps of a representative placenta for uncompensated and fluence-compensated photoacoustic data. Qualitatively, the fluence-compensated data exhibits greater signal amplitude in deeper regions of the placenta than the uncompensated data. The vascular system of the placenta is expected to have highly oxygenated edges near the maternal spiral arteries (decidua) and the umbilical cord (labyrinth), with a dip in the center (junctional zone)[103]. Fig. 7E-F depicts the variations between the distributions of HbT and StO_2_ in compensated and uncompensated volumes. The curves of the HbT distributions have differing shapes, with the non-compensated data having an exponential distribution, while the compensated data displays a right-skewed curve. A two-sided Wilcoxon rank sum test indicates that the medians of the two groups differ significantly (p < 0.0001, z = − 405.53). Comparing the StO_2_ distributions of compensated and uncompensated placenta data using an unpaired two-tailed t-test reveals a statistically significant difference between the means of the two distributions. (p < 0.0001, t = −176.06). While both the uncompensated and compensated distributions have similar shapes, the non-compensated StO_2_ data has a left shifted peak and a variance 1.1542 times higher than the fluence compensated data. Due to the depletion of hemoglobin signal strength at depth, one can see that a significant amount of data is lost in the lower regions of the non-compensated placental data. The fluence compensated data also show higher StO_2_ values in the labyrinth of the placenta, where the active transport of oxygen occurs between mother and fetus.

**Figure 7.**
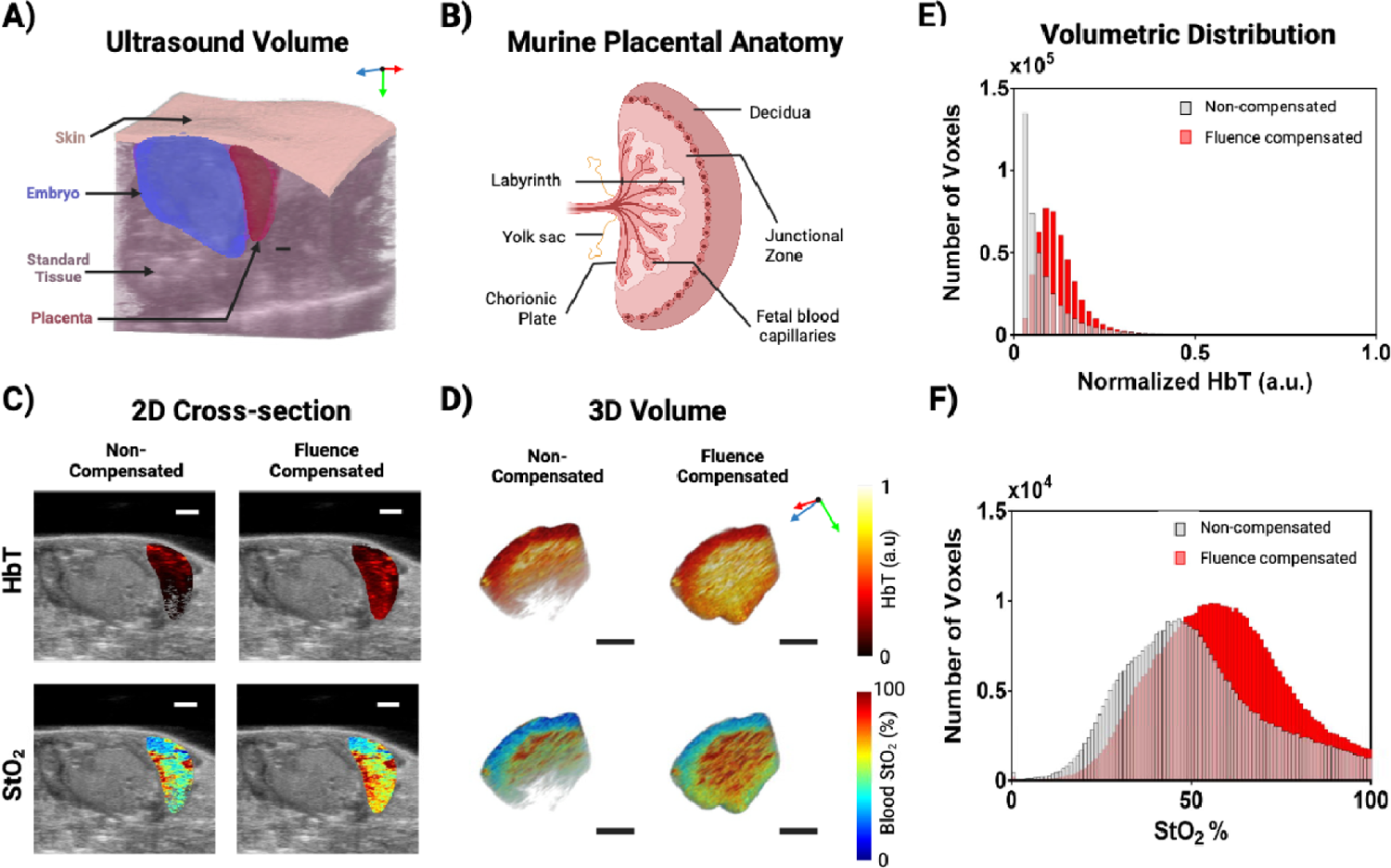
A) 3D ultrasound rendering displaying the tissue region labels used to assign optical properties to the volume. B) Murine placental anatomy displaying the relative locations of the structures in the placenta. C) 2D renderings of placental HbT (top) and StO_2_ (bottom) for non-compensated (left) and fluence compensated (right) data. D) 3D renderings of placental HbT (top) and StO_2_ (bottom) for non-compensated (left) and fluence compensated (right) data. X, Y, and Z-direction are shown with red, green, and blue arrows respectively, where the Z-direction is the transducer scanning direction. All scale bars shown represent 2 mm. E-F) Bar graphs displaying the voxel-wise distribution of placenta HbT (E) and StO_2_ (F) for the placenta rendered in panel A. The non-compensated values are displayed in gray, while the fluence compensated values are displayed in red.

The placenta is a highly vascularized organ and the development and changes in the vascularity, perfusion, and oxygenation provides great insight into the health of the pregnancy. Specifically, vascularity and oxygenation changes can be observed with optical imaging, however it is known that deep imaging of solid vascular systems such as the liver or placenta leads an incomplete image of the vascularity and function due to high light attenuation. Early placental development, placentation, is comprised of angiogenesis and hypoxic conditions[104]. Changes in hemoglobin concentration can detect anemic conditions which give insight on angiogenesis[105], while hypoxia is indicative of vascular remodeling, hypertension, and other stressors[106]. Placental positions in pre-clinical models can vary from superficial and parallel to the transducer to hidden by adipose tissue or the embryo and perpendicular to the transducer, which can create inconsistent StO_2_ and HbT scans and analysis. These inconsistencies do impede precise measurements and comparisons, which then impact understanding and grading the health of different placental disorders. The placenta shown in this work was relatively superficial and unobstructed, and yet fluence compensation still significantly changed the measured StO_2_ and HbT of the placental volume. These results agree with similar results in the field [60, 107]. Fluence compensation can allow more accurate access to information across the entire placenta *in utero* and at earlier time points when the placentas may be more hidden under vital maternal organs. The PHANTOM toolkit will enable researchers to effortlessly segment the placenta, setup the optical properties and depth compensate the photoacoustic signal.

## 4. Conclusions

In this work we have developed an easy-to-use workflow and toolkit named PHotoacoustic ANnotation TOolkit for MATLAB (PHANTOM) for compensating PA images for depth dependent changes in fluence utilizing structural information gathered from US. This toolkit supports users in the segmentation of ultrasound images into 3D labeled tissue structures that may be assigned default or custom optical properties. In addition, we have produced a light source configuration in MCXLAB that replicates the optical fiber illumination used by the VS system. We have displayed the improved accuracy of StO_2_ measurements calculated from this fluence-compensated data in phantom experiments with varying states of blood oxygenation and tissue thickness. The workflow was also tested on two types of *in vivo* data, a subcutaneous tumor model and a placental model of ectopic calcification. For both cases, there were significant differences between in the StO_2_ and HbT volumetric distributions when comparing the non-compensated and fluence compensated data, particularly at deeper depths and validated with immunohistochemistry.

As previously stated, the method of using MC simulations of light propagation can never be perfect, due to the inverse nature of the problem. This issue is exasperated by the limited availability of optical property values in literature, and differences in measurement techniques and wavelengths used for measuring these properties. Due to these reasons, this default optical properties presented in this paper rely heavily on the work of Jacques[20], which provides the most comprehensive method of defining unknown optical properties of tissues. This work however is a decade old and there is a need for an updated compilation of optical property values and the relative blood, water, fat, and melanin volume fractions of different tissue types. In our work, tissue regions are assigned one set of optical properties, whereas actual *in vivo* tissues have a heterogeneous distribution of chromophores that cannot be truly represented by singular values.

Constraints on the reliability of this fluence compensation method must also be considered in relation to image quality. One such factor that must be considered is near-field interference caused by signals generated on the transducer surface when the transducer is situated too close to the tissue boundary. These artifacts can also cause secondary reflections deeper within the tissue. Understanding noise levels has also become increasingly important in PAI. As fluence correction rescales the data, it is probable that the image’s fundamental signal-to-noise ratio (SNR) will not improve, resulting in increased background noise in deep regions. When applying fluence compensation, it is important that one is weary of system and sample dependent noise as these factors will change on a case-by-case basis. As light attenuation decreases the SNR of the image with depth, it is important to understand the noise levels in different areas of an image and use discretion when applying fluence compensation to deeper depths. Rich et al. found that for a 21 MHz transducer similar to the one used in this work had a depth limit of 15 mm to detect signal over noise [108]. The Rose criteria is a widely accepted standard in the field of object detection. According to this standard, a signal must have a magnitude that is at least five times higher than the standard deviation of the background in order to be distinguished from noise [109]. The depth at which fluence compensation will hold is dependent on the wavelength of light, tissue optical properties, number of averages taken, and system-based characteristics such as resolution and gain. Despite this, we clearly demonstrate the improved estimations of blood StO_2_ gathered with PAI by utilizing the MC method of fluence compensation. This was shown for phantom images using real-time pO_2_ measurements as well as subcutaneous tumor images using IF staining.

In this research our methodology was specifically designed for the commercially available VevoLAZR-X system-based fiber delivery systems, the workflow is applicable to any combined ultrasound and photoacoustic imaging system and fiber configuration. Here we also use wavelengths of 750 and 850 nm to calculate StO_2_, however the default optical properties provided with PHANTOM range from 300 - 1000 nm, allowing the toolkit to assist a wider range of applications. The toolkit is adaptable to other endogenous and exogenous photoacoustic contrast agents where the optical properties can be defined within the GUI. All code used to generate the data and simulations discussed in this manuscript are available upon request. PHANTOM will prompt and guide the user through the otherwise complicated and time-consuming task of annotating and fluence compensating large photoacoustic datasets and will be particularly useful for researchers unfamiliar with laser-tissue interactions and outside the field of photoacoustic imaging.

## Acknowledgements

The authors would like to acknowledge Dr. Mary Wallingford for providing the Slc20a2 ectopic calcification mouse model. The authors would also like to acknowledge the members of the integrated Biofunctional Imaging and Therapeutics laboratory, specifically Christopher D. Nguyen, Deeksha Sankepalle, Marvin Xavierselvan, and Vicky Yang for their useful discussions and support. Funding support from National Institutes of Health grants (S10OD026844, R21CA263694 and R01CA231606).

