## Supplementary Data for "Ultrasound-guided Photoacoustic image Annotation Toolkit in MATLAB (PHANTOM) for preclinical applications"

Supplementary Material

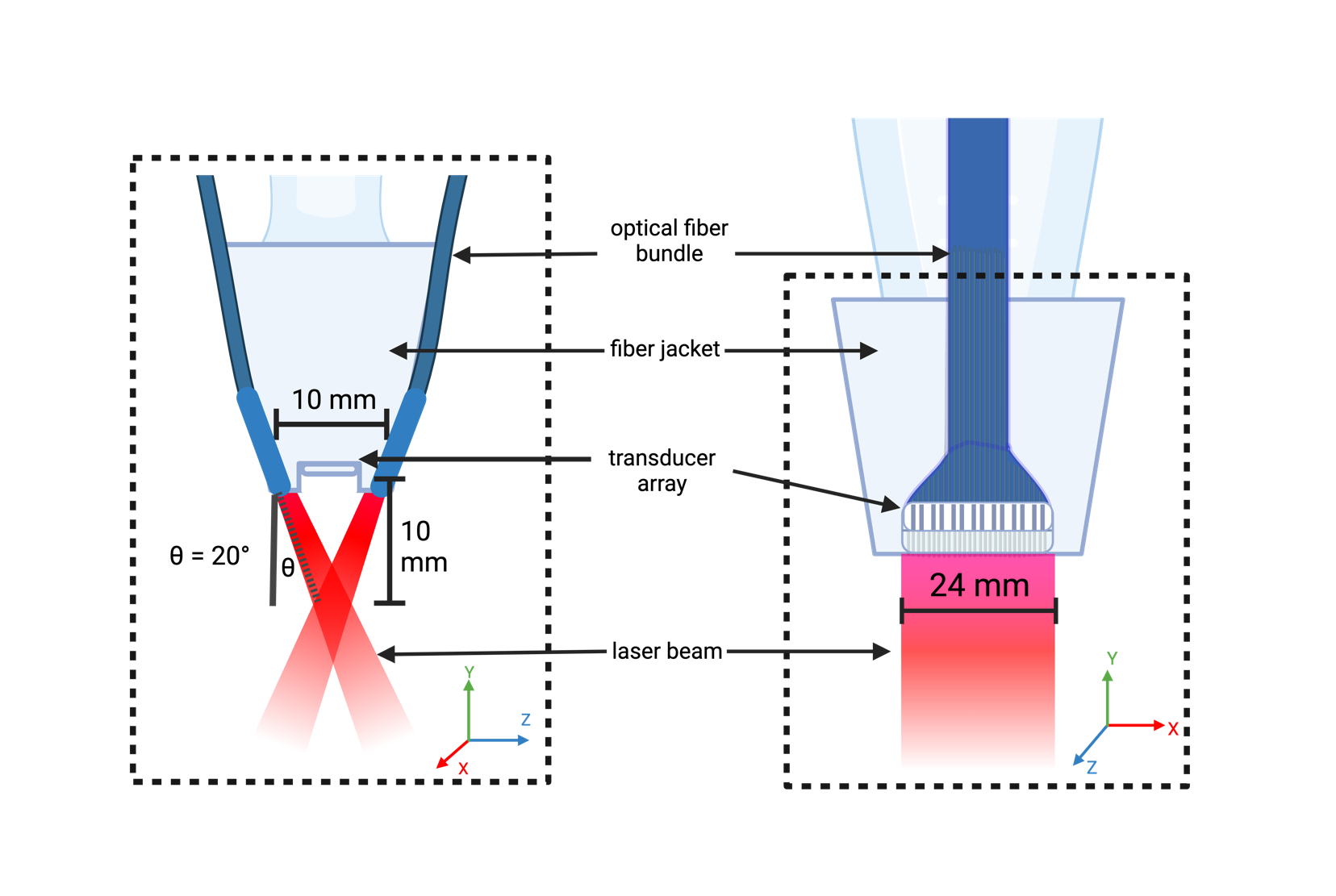

**Figure S1**. Schematic of the Vevo LAZR-X transducer and optical fiber setup for the MX250S transducer paired with the blue optical fibers. X, Y, and Z-direction are shown with red, green, and blue arrows respectively, where the Z-direction represents the transducer scanning direction

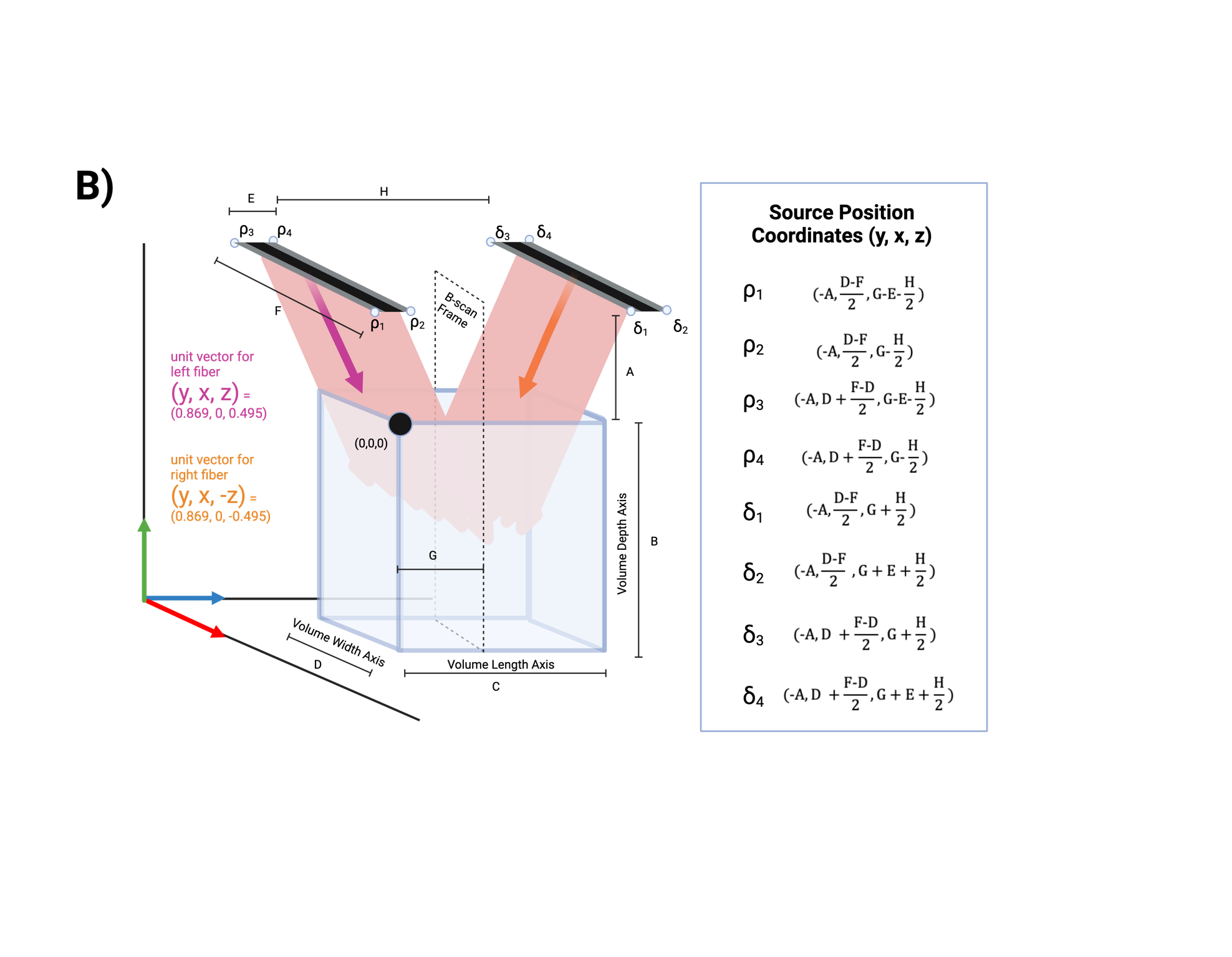

**Figure S2.** Geometry of the light source configuration of the bifurcated optical fibers in relation to the imaging volume and transducer position. . X, Y, and Z-direction are shown with red, green, and blue arrows respectively, where the Z-direction represents the transducer scanning direction

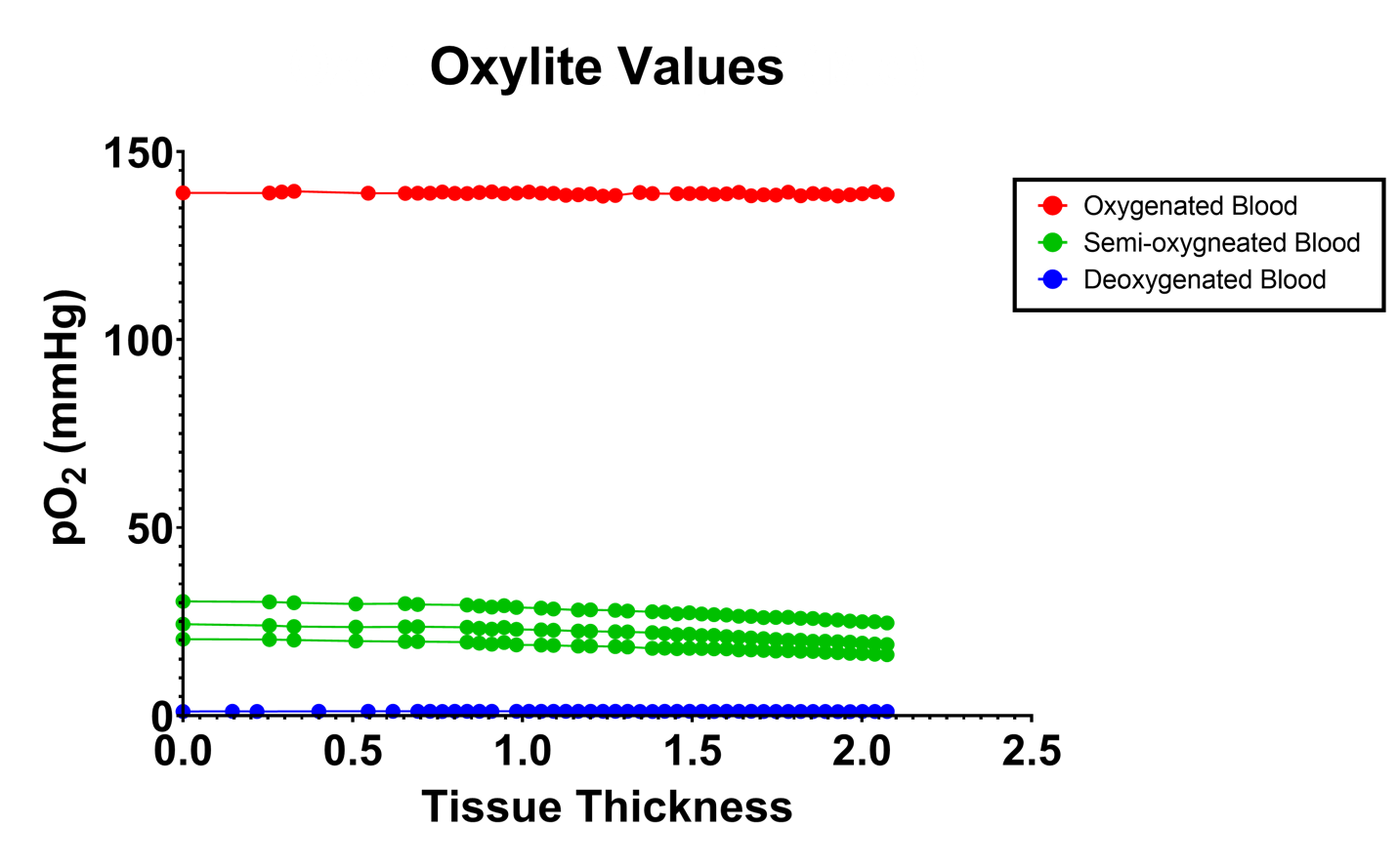

**Figure S3.** Plots of pO2 measured by an oxygen monitor during phantom imaging. Oxygenated blood is shown in red (n=3), deoxygenated blood is shown in blue (n=3), and semi-oxygenated blood is shown in green (n=1 for each line).

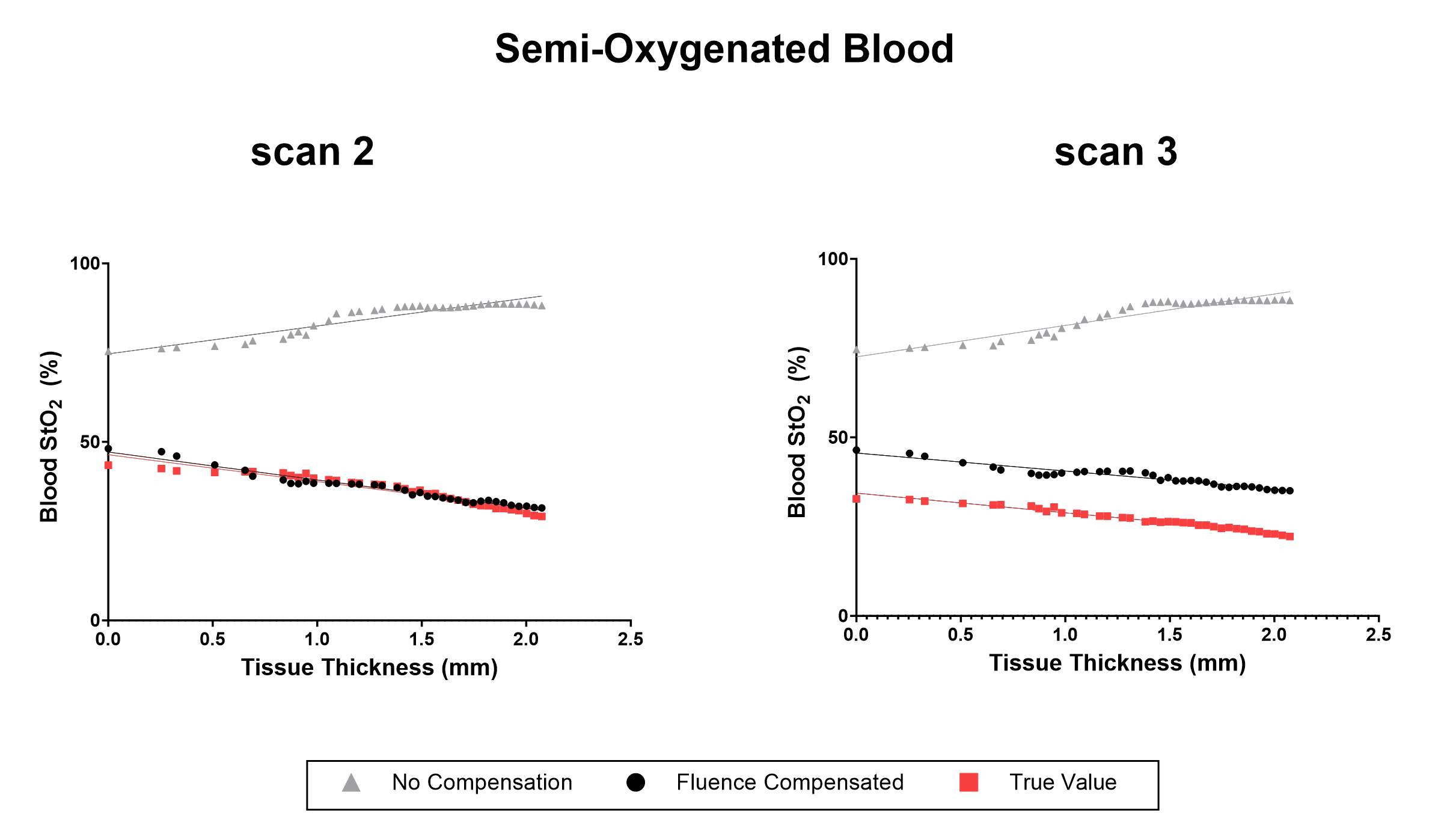

**Figure S4.** Plots of StO_2_ against the thickness of the chicken breast inclusion from the phantom experiment for scans 2-3 of the semi-oxygenated blood condition. The fluence compensated data (FC) is shown in black, the non-compensated data (NC) is shown in gray, while the ground truth values (GT) are plotted in red. The linear fit for the NC, FC, and GT data are also plotted for scan 2 (R^2^ = 0.950, 0.948, 0.860 for GT, FC, and NC respectively) and scan 3 (R^2^ = 0.950, 0.948, 0.860 for GT, FC, and NC respectively)

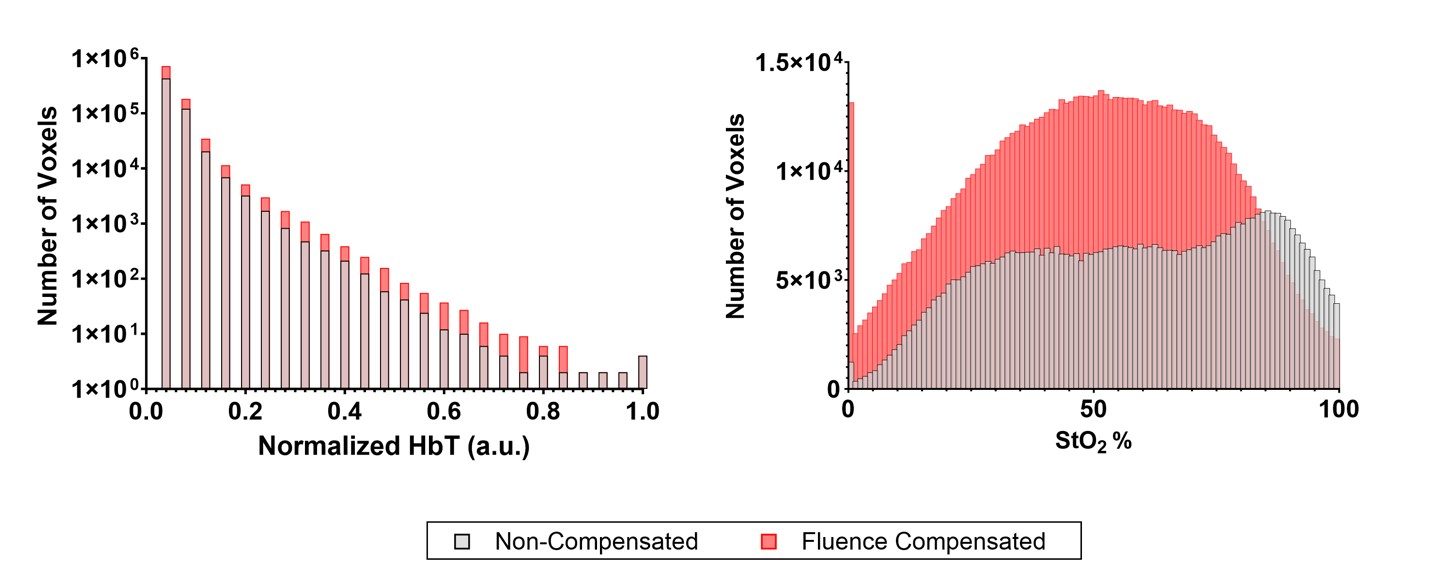

**Figure S5.** Bar graph displaying the voxel-wise distribution of HbT (left) and StO_2_ (right) for the subcutaneous tumor shown in Figure 6 of the main text. The non-compensated values are displayed in gray, while the fluence compensated values are displayed in red.

| **Tissue Type** | **B (blood volume fraction)** | **S (blood oxygen saturation)** | **W (water content)** | **M (melanin volume fraction)** | **F (fat volume fraction)** | **Reference** |
| --- | --- | --- | --- | --- | --- | --- |
| Blood | 1 | Varies (set by user) | 0.95 | 0 | 0 | Jacques^1^ |
| Skin Type I-II | 0.34 | 0.985 | 0.214 | 0.0087 | 0.277 | Jacques^1^  Tseng^2^ |
| Skin Type III-IV | 0.41 | 0.992 | 0.261 | 0.0115 | 0.225 | Jacques^1^  Tseng^2^ |
| Skin Type V-VI | 0.12 | 0.993 | 0.166 | 0.0165 | 0.187 | Jacques^1^  Tseng^2^ |
| Brain | 0.0305 | 0.59 | 0.78 | 0 | 0.6 | O’sullivan^3^  Keep^4^  Chang^5^ |
| Generic Solid Tumor | 0.01 | 0.65 | 0.25 | 0 | 0 | Flexman^6^ |
| Bone | 0.0005 | 0.875 | 0.3 | 0 | 0 | Harrison^7^  Mohammed^8^ |
| Muscle | 0.012 | 0.73 | 0.76 | 0 | 0.2 | Hindel^9^  Miranda-Fuentes^10^  Lorenzo^11^  Marty^12^ |
| Fatty Tissue | 0.05 | 0.7 | 0.15 | 0 | 0.8 | Meglinski^13^  Abe^14^ |
| Liver | 0.495 | 0.767 | 0.773 | 0 | 0.11 | Weaver^15^  Kitai^16^  Lee^17^  Nilsson^18^ |
| Breast (non-cancerous) | 0.0098 | 0.6802 | 0.1602 | 0 | 0.6233 | Jacques^1^ |
| Soft Tissue | 0.54 | 0.76 | 0.11 | 0 | 0.69 | Jakubowski^19^ |
| **Table S1.** Relative chromophore contributions of each tissue type used to calculate absorption coefficient. | | | | | | |

| **Tissue Type** | $\boldsymbol{\alpha'}$ **(mm^-1^)** | $\boldsymbol{f}_{\boldsymbol{Ray}}$ | $\boldsymbol{b}_{\boldsymbol{Mie}}$ | **Reference** |
| --- | --- | --- | --- | --- |
| Blood | 2.2 | 0 | 1.0 | Alexandrakis^20^ Friebel^21^ |
| Skin (all types) | 4.8 | 0.409 | 0.702 | Jacques^1^ |
| Solid Tumor | 3.73 | 0.72 | 0 | Sandell and Zhu^22^ Jacques^1^ |
| Bone | 1.53 | 0.022 | 0.326 | Jacques^1^ |
| Muscle | 1.47 | 0 | 0.926 | Jacques^1^ |
| Fatty Tissue | 1.93 | 0.174 | 0.447 | Jacques^1^ |
| Liver | 1.65 | 0 | 1.64 | Yi and Backman^23^ |
| Breast (non-cancerous) | 1.87 | 0.288 | 0.685 | Jacques^1^ |
| Soft Tissue | 1.91 | 0.153 | 1.091 | Jacques^1^ |
| Brain | 2.74 | 0.315 | 1.087 | Jacques^1^ |
| **Table S2.** Parameters of $\alpha$’, f_ray,_ and b_Mie_ of each tissue type used to calculate scattering coefficient. | | | | |
